# Cycle-by-cycle respiration waveforms are coupled with the shape of neural oscillations

**DOI:** 10.64898/2026.04.13.718339

**Authors:** Eena Kosik-Rose, Guangyu Zhou, Andrew Sheriff, Joshua M. Rosenow, Stephan U. Schuele, Chima O. Oluigbo, Saige Anabel Teti, Mohamad Koubeissi, Md Rakibul Mowla, Ariane E. Rhone, Sukhbinder Kumar, Brian Dlouhy, Christina Zelano, Bradley Voytek

## Abstract

Beyond sustaining life, breathing is a vital physiological rhythm that shapes cognition, perception, emotional regulation, and mental health. Breathing has a direct effect on neuronal excitability and is coupled to neural oscillations across a variety of brain regions. Notably, both respiration and neural oscillations are asymmetric and not perfectly rhythmic: for example, every breath has a different shape and duration, and is interspersed with variable pauses. Here, we examined the coupling between breathing and the brain by quantifying the nonsinusoidal features of each breath and comparing it to the shape of each corresponding neural oscillation cycle. By leveraging invasive human brain recordings from 16 participants, we found respiration-neural waveform coupling on a breath-by-breath, cycle-by-cycle basis across limbic and cortical forebrain regions. For decades, the dominant perspective on cognition and mental health have focused on the brain, but recent work is highlighting the importance of brain-body interactions. Our results show that the coupling between breathing and neural activity is much richer than previously appreciated, and our approach opens new avenues for studying these peripheral-to-central nervous system interactions in a more robust, temporally precise manner.

## Introduction

Breathing is a vital rhythm of life, sustaining metabolic and homeostatic processes. While the brainstem mechanisms related to the control of breathing have long been understood, more recent work has shown coordinated dynamics between respiration and higher-order cognitive and affective processes as well (Allen et al., 2021; Brændholt et al., 2023; Del Negro et al., 2018; Herrero et al., 2018; Kluger & Gross, 2021). While the mechanisms that relate respiration to cognitive and affective processes are less understood, in several brain areas, neural responses to various external and internal stimuli have been shown to be dependent on the phase of respiration cycle (Brændholt et al., 2025; Harting et al., 2025; Mizuhara & Nittono, 2023; Park et al., 2020). Behaviorally, this manifests in perceptual, memory, and emotion-related tasks, where participants show significantly different performance depending on if a stimulus appears during the inhalation or exhalation phase of the breath (Arshamian et al., 2018; Grund et al., 2021; Johannknecht & Kayser, 2022; Kluger & Gross, 2020, 2021; Nakamura et al., 2022; Perl et al., 2019; Zelano et al., 2016). Notably, humans and other animals breathe asymmetrically with varying inhale and exhale durations. This results in a behavioral asymmetry such that perception and stimulus detection are faster and more accurate during the transition between inhalation and exhalation – approaching the peak of the airflow (Grund et al., 2021; Zelano et al., 2016).

One mechanistic hypothesis for this behavioral asymmetry is breathing’s apparent influence on neural local field potentials (LFPs). Respiration can entrain LFPs in the mammalian forebrain, particularly in regions implicated in sensory, cognitive, and affective function (Herrero et al., 2018; Kluger & Gross, 2021; Zelano et al., 2016). This respiration-entrained rhythm appears to be linked to the nasal-specific breathing pathway. When the nasal pathway is bypassed, whether via tracheostomy in rats or mouth breathing in humans, respiration-locked brain rhythms disappear (Fontanini et al., 2003; Ito et al., 2014; Zelano et al., 2016). However, emerging research has called this into question in humans, suggesting multiple afferent pathways for this neural synchronization, such as from the trachea, lungs, diaphragm, and chest wall (Dlouhy et al., 2025). Respiration’s influence on neural activity has even been extended to modulation of single-unit membrane potential and spiking discharge in these non-olfactory regions (De Falco et al., 2024; Juventin et al., 2022; Karalis & Sirota, 2022).

Work assessing the relation between respiration and neural activity has predominantly relied on the metric of coherence to examine respiration-brain synchronization, leveraging the frequency-specific consistency in phase relationships between these signals (Tort, Brankačk, et al., 2018; Tort et al., 2025; Tort, Ponsel, et al., 2018). While informative and an important first step, coherence summarizes coupling over a longer time scale because it averages over multiple cycles and does not capture the fine-grained temporal coupling of individual respiratory and neural cycles. This is particularly important given the substantial variability in individual breath morphology – including asymmetries in rise and decay durations, differences in inhale peak volume, and inter-breath pauses – which can be masked using traditional approaches that assume respiration is a stable rhythm (Noto et al., 2018). This heterogeneity in oscillatory waveform shape has been clearly demonstrated in LFP signals, which have long been known to be asymmetric and bursty (Bender et al., 2025; S. R. Cole & Voytek, 2017; S. Cole & Voytek, 2019; Jones, 2016; Trimper et al., 2014). These nonsinusoidal features in the LFPs have been linked to a wide variety of cognitive and disease states, ranging from memory performance in rats (Trimper et al., 2014) to the efficacy of deep brain stimulation in people with Parkinson’s disease (S. R. Cole et al., 2017; Jackson et al., 2019). Whether respiration and neural oscillations are coupled not just in frequency, but in the fine-grained shape of each cycle, on a breath-by-breath basis, remains unknown.

In this study, we explicitly sought to examine the links between respiration and neural activity by leveraging the nonsinusoidal and heterogeneous nature of both signals. Specifically, we examine whether respiration and the neural LFP exhibit systematic, cycle-by-cycle alignment in waveform morphology. To do this, we introduce a novel, time-domain, cycle-resolved analysis that quantifies waveform shape features of both the respiration signal and LFP on a paired-cycle basis. We analyzed three human intracranial EEG (iEEG) datasets to quantify respiration waveform shape and its relationship to the shape of respiration-entrained neural oscillations across the forebrain at rest. We demonstrate that significant coherence between respiration and neural rhythms, while informative, does not inherently capture the cycle-by-cycle waveform dynamics between these signals. We provide, for the first time, strong evidence that the shape of every breath is coupled with the shape of the LFP on a breath-by-breath, cycle-by-cycle basis. Our results provide compelling evidence of respiration’s fundamental role in modulating neural activity.

## Methods

### sEEG and respiration datasets

We analyzed previously collected data from three different recording locations. Sixteen participants with medically intractable epilepsy (10 participants from Northwestern Memorial Hospital, 3 participants from Children’s National Hospital, and 3 participants from University of Iowa Stead Family Children’s Hospital) underwent stereoelectroencephalography (sEEG) during a 2-week monitoring period for seizure focus localization. Participants’ demographic data are shown in **Table 1**. All participants gave written informed consent and all methods were approved by the Institutional Review Board of the respective institutions. Participants and/or guardians could withdraw consent or assent at any point without affecting their clinical care. Electrode placement and respiratory monitoring techniques did not deviate from standard clinical procedures at each institution. All data ranged from 1.5 to 15.5 minutes.

**Table 1:** Clinical and demographic information for all participants included in the study. Participants were recruited from three sites: Northwestern University (participants 1–10), Children’s National Hospital (participants 11–13) and the University of Iowa Stead Family Children’s Hospital (participants 14–16). All participants were undergoing intracranial EEG monitoring for surgical evaluation of treatment-resistant epilepsy.

| Participant | Site | Gender | Age at Surgery | Handedness | Duration of Epilepsy (years) | Epileptogenic Zone | Brain MRI |
| --- | --- | --- | --- | --- | --- | --- | --- |
| 1 | Northwestern | Male | 32 | Left | 9 | Left temporal lobe | Normal |
| 2 | Northwestern | Female | 29 | Right | 7 | Left temporal lobe | Normal |
| 3 | Northwestern | Male | 49 | Right | 26 | Left temporal lobe | Chronic stroke/encephalomalacia in left putamen, insula, parietal cortex |
| 4 | Northwestern | Male | 48 | R | 11 | Right temporal lobe | Normal |
| 5 | Northwestern | Male | 32 | Right | 10 | Left basal temporal | Normal |
| 6 | Northwestern | Female | 27 | Right | 5 | Left mesial temporal | Left mesial temporal sclerosis |
| 7 | Northwestern | Male | 54 | Left | 2 | Right mesial temporal | Normal |
| 8 | Northwestern | Female | 25 | Right | 3 | Left mesial temporal | Normal |
| 9 | Northwestern | Female | 50 | Right | 27 | Left temporal | Heterotopic gray matter to left atrial ependyma |
| 10 | Northwestern | Female | 36 | Right | 4 | Left frontal lobe | Possible moderate mesial temporal sclerosis |
| 11 | CNMC | Male | 19 | Right | 18 | Right temporal | Right amygdalohippocampectomy |
| 12 | CNMC | Female | 19 | Right | Unknown | Left temporal | Left amygdalectomy |
| 13 | CNMC | Male | 21 | Right | 6 | Focal, right frontal | Possible dysgenesis/dysplasia of right hippocampus |
| 14 | Iowa | Female | 12 | Right | 8 | Multifocal, but right mesial temporal preponderance | Normal |
| 15 | Iowa | Male | 16 | Right | 9 | Left anterior temporal lobe; mesial temporal lobe | Hypometabolism in the left media and anterior temporal lobe |
| 16 | Iowa | Female | 14 | Right | 10 | Right parieto-temporal region | Prior laser ablation of right posterior temporoparietal cortical dysplasia |

For participants at Northwestern Memorial Hospital and Children’s National Hospital, the sampling rate, which was determined clinically, was 1,000 Hz or 2,000 Hz for all participants. A surgically implanted electrode strip facing towards the scalp or a surface electrode served as the reference and ground. Respiratory signals were recorded using a piezoelectric pressure transducer (Salter Labs BiNAPs 5500) attached to a nasal cannula placed at the participant’s nose, and a breathing belt placed around the participant’s abdomen (Perfect Fit II Adult Effort Belt, DyMedix Diagnostics). Participants were instructed to sit comfortably breathing through their noses with their eyes open while sEEG data were recorded using a clinical Nihon Kohden acquisition system with a built-in high-pass filter at 0.008 Hz. For participants at University of Iowa, they were implanted with intracranial electrodes (DIXI Medical US Products) at sites determined by the multidisciplinary epilepsy team. The sampling rate was 2000 Hz, and the filter setting at acquisition was 0.1 Hz to 500 Hz. A single subgaleal depth electrode was used as the reference. Respiratory signals were recorded using a piezoelectric pressure transducer (Salter Labs BiNAPs 5500) attached to a nasal cannula placed at the participant’s nose, and a chest/abdominal plethysmography belt (ProTech zRIP). Participants were instructed to rest quietly in the hospital bed while sEEG data were recorded to a Neuralynx ATLAS Neurophysiology System (Boseman, MT). Across all sites, 10 participants had respiration recordings from both modalities, 3 had only nasal cannula, and 3 had only breathing belt.

### Data preprocessing

LFP and respiratory data were downsampled to 500 Hz using MNE in Python (version 3.11.10) (Gramfort, 2013). Recordings were bipolar re-referenced by subtracting the activity of adjacent electrode contact pairs to reduce noise, eliminate far-field effects, and decrease concerns over intracranial CSF pulsations and electrode movement synchronized to respiration. Respiratory signals were further low-pass filtered at 2 Hz using a finite impulse response (FIR) filter as implemented in MNE.

To determine the electrode locations for participants at Northwestern Memorial Hospital, the following procedure was used, as described in Zhou et al., 2021. The postoperative computed tomography images were registered to individual preoperative structural MRI images using FSL’s linear registration tool (Jenkinson et al., 2002; Jenkinson & Smith, 2001). The transformed post-operative computed tomography images were thresholded and each electrode was identified manually. We used the center of each identified electrode as its coordinate in individual anatomical space. In order to visually represent electrodes across participants in one image, electrode coordinates were converted into MNI space by normalizing individual MRI images to a standard MNI brain (MNI152_1mm_brain) using FSL. We also performed subcortical segmentation of each individual’s MRI image using FreeSurfer (Fischl, 2012; Fischl et al., 2002; RRID:SCR_001847).

For the participants at the Iowa Stead Family Children’s Hospital, the following procedure was used, as described in Harmata et al., 2023. Whole-brain high-resolution T1-weighted structural MR scans (resolution and slice thickness ≤ 1.0 mm) were obtained before and after electrode implantation. Post-implantation CT and 3T MR structural scans were linearly coregistered to pre-implantation MR scans using the FLIRT module of FSL (Jenkinson et al., 2002). These images were corrected manually for displacement and deformation from surgery using nonlinear thin-plate spline warping using manually selected control points. Finally, FreeSurfer was used to identify the cortical surface and white/gray boundaries within pre-implantation T1 images, which were corrected manually based on visual inspection where necessary.

Raw region labels, which varied in nomenclature across atlases and sites, were harmonized to a common set of anatomical categories using a custom label-normalization procedure applied uniformly across all participants. The atlas column in the electrode data frame records the source atlas for each participant. Electrode coverage by anatomical region for both the airflow and belt datasets is summarized in **Table 2**.

**Table 2:** Electrode coverage. Number of electrodes and participants contributing to each anatomical region for both the airflow and belt datasets.

|  | **Airflow** |  | **Belt** |  |
| --- | --- | --- | --- | --- |
|  | **No. of Electrodes** | **No. of Participants** | **No. of Electrodes** | **No. of Participants** |
| Amygdala | 37 | 11 | 46 | 12 |
| Hippocampus | 63 | 11 | 68 | 12 |
| Parahippocampal | 12 | 4 | 11 | 5 |
| Insula | 86 | 10 | 66 | 10 |
| Orbitofrontal cortex | 53 | 7 | 61 | 8 |
| Inferior frontal gyrus | 62 | 10 | 49 | 10 |
| Frontal–rostral middle | 36 | 6 | 21 | 5 |
| Frontal–caudal middle | 25 | 5 | 24 | 4 |
| Frontal–superior | 36 | 5 | 29 | 3 |
| Precentral | 58 | 5 | 67 | 6 |
| Paracentral | 5 | 1 | 5 | 1 |
| Postcentral | 17 | 7 | 20 | 8 |
| Cingulate cortex | 33 | 4 | 35 | 4 |
| Parietal–inferior | 30 | 4 | 30 | 4 |
| Parietal–superior | 14 | 2 | 13 | 1 |
| Occipital cortex | 9 | 5 | 7 | 4 |
| Fusiform | 25 | 8 | 32 | 10 |
| Temporal–middle | 104 | 13 | 126 | 13 |
| Temporal–inferior | 36 | 11 | 50 | 12 |
| Temporal– | 88 | 12 | 93 | 13 |
| superior |  |  |  |  |
| Supramarginal | 63 | 8 | 66 | 9 |
| Transverse<br>temporal | 9 | 2 | 8 | 1 |
| Thalamus | 36 | 4 | 29 | 4 |
| White matter | 503 | 13 | 386 | 13 |
| CSF | 19 | 6 | 19 | 6 |
| Total | 1459 | 13 | 1361 | 13 |

### Coherence

To calculate coherence between the LFP and the breathing signals, we estimated cross-spectral and auto-spectral densities using the magnitude-squared coherence function implemented in ‘scipy.signal.coherence’ (Virtanen et al., 2020). Signals were segmented into epochs of 32 s with an overlap of 27 s following parameters from Herrero et al., (2018), where the 32 s window length provides a frequency resolution of 0.03 Hz (1/32 Hz), sufficient to capture the slow respiration rhythm. The magnitude-squared coherence was calculated as:

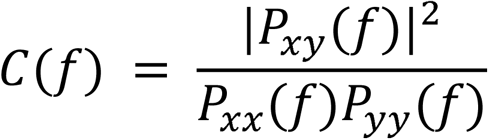

where

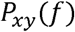

is the cross-spectral density and

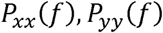

are the auto-spectral densities of the neural and respiratory signals, respectively.

For each neural channel, the maximum coherence value within each individual’s breathing frequency range was extracted. This frequency range was determined by first computing the respiratory power spectral density (PSD) via Welch’s method (Welch, 1967) using a 40 s Hanning window with a 20 s overlap. This long window is used to provide adequate frequency resolution (∼0.025 Hz) to account for the slow respiration rhythm. Then, using the *specparam* toolbox (Donoghue et al., 2020), the center frequency of the highest amplitude peak was defined as the breathing frequency. A frequency band (0.2 Hz bandwidth) centered at the breathing frequency was used to bound the expected respiration-related frequency range in the coherence analysis.

To assess the statistical significance of coherence, we generated 1,000 surrogate neural signals for each LFP channel through phase randomization. To achieve this, LFP data were transformed to the frequency domain via Fast Fourier Transform (FFT), their phases were randomized while preserving amplitude spectra, and the LFPs were reconstructed in the time domain with inverse FFT. The coherence between each surrogate and the respective participant’s respiratory signal was calculated, and a critical threshold was defined as the 99th percentile of surrogate coherence values. Coherence exceeding the threshold within the breathing frequency range was considered statistically significant (*p* < 0.01). Channels were retained for the next stage of analysis only if coherence at the respiration-frequency peak exceeded the surrogate-derived threshold and the neural PSD had a *specparam*-identified peak within the same ± 0.1 Hz breathing frequency band. This dual criterion was applied based on the recommendation from Tort et al., (2025). It is possible that significant coherence and a local PSD peak can dissociate – a channel may show coherence without a local power peak or exhibit high coherence driven by harmonics of the breathing frequency without a fundamental peak due to the nonsinusoidal waveform morphology.

### Cross-correlation analysis

Following the coherence-based selection of neural channels, we applied the first of two time-domain screening steps to ensure that the subsequent waveform shape analysis was appropriate: cross-correlations within fixed windows centered on the respiration peaks to provide consistent alignment across cycles.

First, neural and respiration signals were bandpass filtered between 0.01 and 2 Hz using a fifth-order Infinite Impulse Response (IIR) Bessel filter applied forward-backward (zero-phase) to preserve time-domain fidelity (Ghibaudo et al., 2023) and subsequently z-scored to normalize across time. Both signals were segmented into epochs centered on each respiration peak (corresponding to peak inhalation), with window lengths calibrated to each participant’s breath cycle. Because windows were sized to the breath cycle, both the inhale peak and exhale trough were captured within each epoch.

Then, we computed the cross-correlation function (CCF) between the neural and respiration signals for each epoch, and then averaged CCFs across epochs yielding a mean CCF per channel, using ‘scipy.signal.correlatè. Lags were restricted to half of the respiration cycle to minimize spurious associations within a cyclical signal. From this restricted window, we extracted the maximum absolute value of the mean CCF for each LFP channel.

Statistical significance of the CCF was assessed using a permutation null distribution in which respiration epochs were paired with randomly sampled neural segments of equal length (drawn outside a 10 s margin around the original window). For each permutation (500 iterations), a mean CCF was calculated from which the maximum absolute CCF value was extracted, yielding a channel-specific null distribution. The significance threshold was defined as the 99th percentile of the null distribution (*p* < 0.01). Channel polarity was flipped at this stage when the mean CCF peak < 0 to account for arbitrary polarity from bipolar re-referencing.

### Respiration phase monotonicity index

As a second time-domain screening step, we quantified whether the neural signals were consistently monotonic across the rising and falling phases of the respiration cycle using what we call a phase monotonicity index (PMI), adapted from the cycle-by-cycle analysis framework from S. Cole & Voytek, (2019).

Respiration cycles were first defined using troughs and peaks identified from the breathing signal. For each cycle, the corresponding neural signal was partitioned into a rising phase (trough to peak) and a falling phase (peak to subsequent trough). Within each phase, we computed the fraction of consecutive sample-to-sample differences that were directionally consistent with monotonic increase or decrease. A channel-level PMI value was obtained as the median PMI across all valid respiration cycles.

Statistical significance of the observed PMI was assessed using a circular-shift permutation null distribution. Neural signals were circularly shifted by a random offset of at least 10 s to disrupt their alignment with the respiration cycle while preserving temporal autocorrelation structure. For each permutation (500 iterations), the PMI statistic was recomputed, yielding a null distribution against which the observed PMI was compared. The significance threshold was defined as the 99th percentile of the null distribution (*p* < 0.01), and channels were retained if the observed PMI exceeded the threshold.

### Calculation of waveform shape features

#### Peak and trough detection

Peaks and troughs were extracted separately for neural and respiration signals using a two-stage adaptive procedure. In the first stage, candidate extrema were identified using ‘scipy.signal.find_peaks’, applying a base height threshold and a minimum inter-peak distance scaled to the sampling rate (distance = 1.5 × sampling frequency (fs)). A second pass refined detection by computing the median peak amplitude and applying a relative amplitude threshold. To ensure physiologically interpretable alternation between peaks and troughs, consecutive extrema of the same type were resolved by retaining only the more extreme point (higher peaks or deeper troughs). Cycles with abnormally large amplitude or root mean square (RMS; indicative of artifact or transient noise) were removed using robust median absolute deviation (MAD) thresholds.

#### Neural-respiration peak matching

To pair neural cycles with their corresponding respiratory cycles, we matched peaks in a one-to-one manner using a greedy nearest-neighbor procedure, which included iterating through each respiration peak in order and assigning it the temporally closest unmatched neural peak. The closest neural peak was accepted as a match if it was delayed less than one respiration period (τ), derived from the dominant respiration frequency. Peak pairs exceeding this threshold or lacking a following trough were excluded.

#### Cycle segmentation and waveform metrics

For each matched peak pair, cycle boundaries were defined using the nearest troughs immediately before and after each peak. Multiple waveform shape features were computed for both respiration and sEEG signals based on previous work (S. Cole & Voytek, 2019; Noto et al., 2018): rise time (onset trough time to peak time), decay time (peak time to subsequent trough time), cycle duration (sum of rise and decay times), amplitude (peak amplitude minus trough amplitude), sharpness (mean absolute first derivative within a full-width-at-half-maximum (FWHM) window around inhale peaks and exhale troughs, normalized by local amplitude), area-under-the-curve (AUC; trapezoidal integration between median-crossing boundaries around inhale peak and exhale trough segments).

All resulting features were stored in a Pandas DataFrame (McKinney, 2010) containing one row per matched neural-respiration cycle.

To reduce the influence of extreme values, we applied feature-wise outlier exclusion. For each waveform metric (rise time, decay time, rise-decay symmetry, cycle duration, inhale peak/exhale trough sharpness and AUC), cycles exceeding 3 standard deviations above the median were discarded.

### Statistical analysis

To characterize the waveform shape features of both respiratory recording modalities, we compared airflow and belt features directly. For each feature, the median value across cycles was computed per participant, and paired differences between modalities were tested using the Wilcoxon signed-rank test across participants with valid recordings from both (N = 10). Because the two streams may differ systematically in their waveform properties, all subsequent analyses were conducted separately for airflow and belt.

Respiratory and neural waveform shape relationships were evaluated using Bayesian multilevel models fit in R v.4.5.2 (R Core Team, 2025) using the *brms* package v.2.23.0 (Bürkner, 2017). For each feature pair, neural features were modeled as a function of matched respiratory features using a within-between decomposition (Hamaker & Muthén, 2020; Mundlak, 1978). The respiratory predictor was split into within-subject and between-subject components, both scaled by their pooled standard deviations (SDs) to standardize coefficients comparable across features.

Compared to frequentist approaches, which determine whether there is an effect, Bayesian models output a probability distribution over the possible values of a parameter, called the posterior distribution. This enables direct probability statements about effect direction and magnitude rather than binary hypothesis testing (McElreath, 2020). Bayesian models were used because they provide stable estimates of complex random-effects structures and naturally accommodate the hierarchical nesting of participants and channels. We chose to use a Student’s t likelihood, instead of a Gaussian, to accommodate any skewed data. Weakly informative priors were placed on all fixed effects (β ∼ Normal(0, 1)), the intercept (Normal(0, 2)), variance components (Exponential(1)), and the degrees-of-freedom parameter (ν ∼ Gamma(2, 0.1)). The Markov chain Monte Carlo (MCMC) algorithm was applied with four chains of 3,000 iterations (1,500 warmup iterations) to obtain the posterior distribution for each parameter. Model adequacy was assessed via posterior predictive checks and convergence determined by the Rhat parameter (the ratio of the between to within chain variance) (Vasishth et al., 2018). For each model, the within-subject slope (β_within), its 95% credible interval (95% CrI), and the probability of β_within being greater than zero, P(β > 0), are reported. Note that credible intervals (CrI) are a Bayesian quantity distinct from frequentist confidence intervals (CI): a 95% CrI directly expresses that, given the data and priors, there is a 95% probability that the parameter lies within the stated range, which is not an interpretation available to frequentist CIs.

To assess whether observed coupling exceeded chance, we fit null models identical in structure to the primary models but using shuffled respiratory predictors. For each feature, the within-subject respiratory predictor (resp_within) was randomly permuted across cycles within each subject, disrupting cycle-by-cycle pairing while preserving the overall distribution of respiratory values and the multilevel structure of the data. Null models were fit using the same priors, sampler settings, and random effects structure as the primary models. Posterior distributions of β_within from null models are shown alongside those from real models in **Fig. 4a**; separation between real and null posteriors provides visual evidence of coupling beyond what would be expected under cycle-label randomization.

## Results

### Respiration waveform shape is heterogeneous and can be rigorously quantified

We first quantified respiration waveform shape on a cycle-by-cycle basis using both nasal airflow and belt recordings (**Fig. 1a**). For each breath, we extracted rise time, decay time, total cycle duration, rise-decay symmetry, inhale peak and exhale trough AUC, and inhale peak and exhale trough sharpness. These features captured the inherent asymmetry and diversity in shape of natural breathing.

**Fig. 1.**
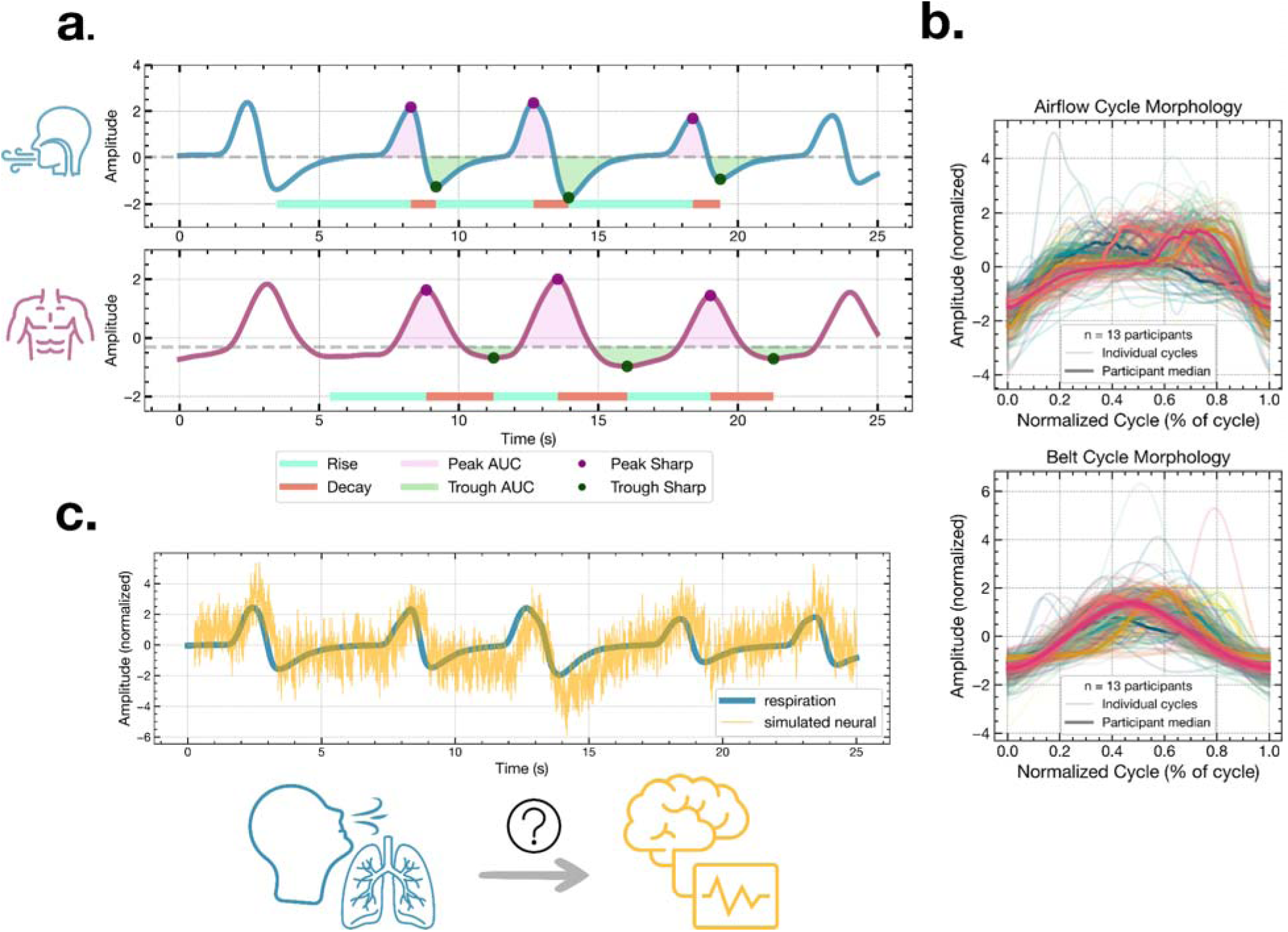
Hypothesis and waveform shape quantification. **a.** Quantification of respiration waveform shape. Airflow (top) and belt (bottom) signals shown with the following waveform features: rise time, decay time, area under the inhale peak, area under the exhale trough, inhale peak sharpness, and exhale trough sharpness. **b**. Overlay of individual respiratory cycles time-normalized to percentage of cycle, shown separately for airflow (top) and belt (bottom). Each colored trace represents an individual cycle; the dark overlaid line shows the cross-cycle mean. The spread of traces illustrates the heterogeneity of waveform morphology both within and across individuals and modalities. **c.** Simulated example illustrating the central hypothesis: a respiration signal (blue) with variable cycle morphology is shown alongside a simulated neural signal (yellow) whose waveform shape tracks that of the respiration cycle-by-cycle. The central question of this paper is whether this waveform morphological coupling is observed in real intracranial EEG.

Respiratory waveform morphology varied significantly across recording modalities (**Fig. 1b**; **Fig. S1**). To formally test modality differences, we compared within-subject airflow versus belt features using paired Wilcoxon signed-rank tests across participants who had both modalities collected (N = 10). These two modalities were significantly different in decay time (*p* = 0.048), inhale peak AUC (*p* = 0.019), exhale trough AUC (*p* = 0.009), exhale trough sharpness (*p* = 0.001), and inhale peak sharpness (*p* = 0.009). In contrast, we didn’t find any difference in rise time (*p* = 0.105), rise-decay symmetry (*p* = 0.064), and cycle duration (*p* = 0.193). The fact that cycle duration did not differ between modalities implies that the airflow and belt yield comparable respiration rates despite systematic differences in waveform morphology. These results confirm that human respiration is nonsinusoidal – as evidenced by the asymmetry between rise and decay times and between inhale and exhale AUC – and exhibits rich, quantifiable morphology that varies meaningfully across cycles and modalities (Noto et al., 2018).

### Respiration-neural coherence and waveform shape coupling identifies widespread respiration-locked activity

We next examined frequency-domain coupling between respiration and neural signals using magnitude-squared coherence within each participant’s dominant breathing frequency band (**Fig. 2a–c**). Across airflow recordings (N = 13 participants; 1,459 channels total) and belt recordings (N = 13 participants; 1,361 channels total), significant respiration-locked coherence (p < 0.01; surrogate-based threshold) was observed in distributed cortical and limbic regions. Channels were retained for waveform shape analysis only if: (i) coherence exceeded the 99th percentile of phase-randomized surrogates within ±0.1 Hz of the respiration peak frequency, and (ii) the neural power spectrum contained a corresponding periodic peak. This procedure identified 104 airflow-coupled channels across 13 participants and 109 belt-coupled channels across 13 participants (**Fig. 3a–b**, left panels).

**Fig. 2.**
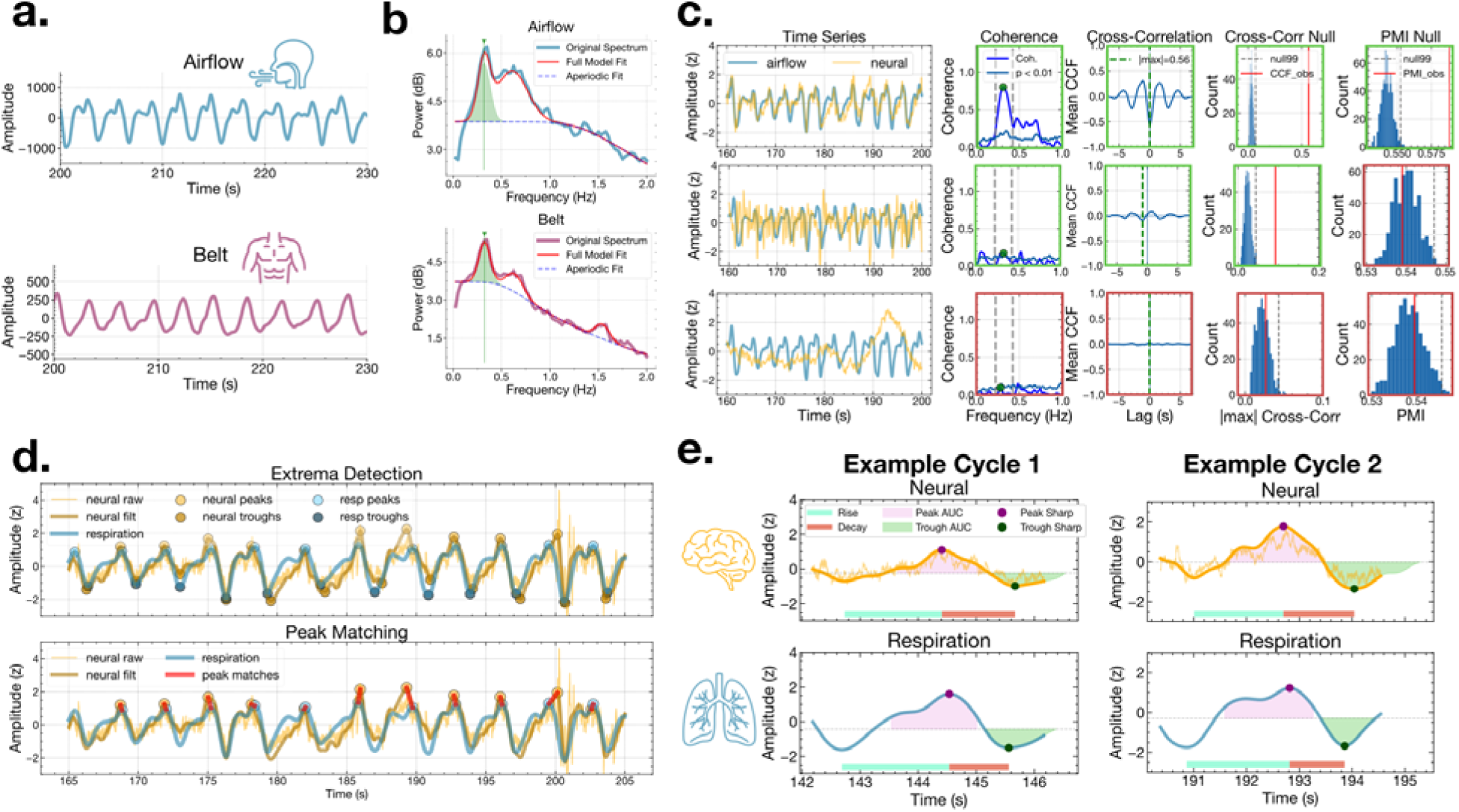
Data analysis pipeline. **a.** Example airflow (top) and belt (bottom) traces from one participant. **b.** Power spectral density of airflow (top) and belt (bottom) signals fit via *specparam*, with the identified respiration peak frequency marked (green arrow). **c.** Three example neural channels (yellow) illustrating the sequential screening criteria for the example airflow channel (light blue). Coherence significance was assessed against 1,000 phase-randomized surrogate channels; CCF and PMI significance were assessed against permutation null distributions (500 iterations each). Only channels passing all three criteria were retained for waveform shape analysis. For each channel (row), columns show the time series (airflow and neural), coherence, mean CCF, CCF null distribution, and PMI null distribution. Green outlines indicate the channel passed that criterion; red outlines indicate failure. Top row: a channel passing all screening criteria; significant coherence, CCF, and PMI exceeding the null. Middle row: a channel with significant coherence and CCF, but PMI failing time-domain screening. Bottom row: a channel failing coherence and all subsequent criteria. **d.** Extrema detection (top) and peak matching (bottom) on filtered and z-scored neural and respiratory signals. Matched peak pairs (red lines) defined the cycles used for feature extraction. **e.** Two example matched neural-respiratory cycle pairs with waveform shape features annotated: rise time (teal), decay time (red), inhale peak AUC (pink shading), exhale trough AUC (green shading), inhale peak sharpness (purple), and exhale trough sharpness (dark green). CCF, cross-correlation function; PMI, phase monotonicity index; AUC, area under curve.

**Fig. 3.**
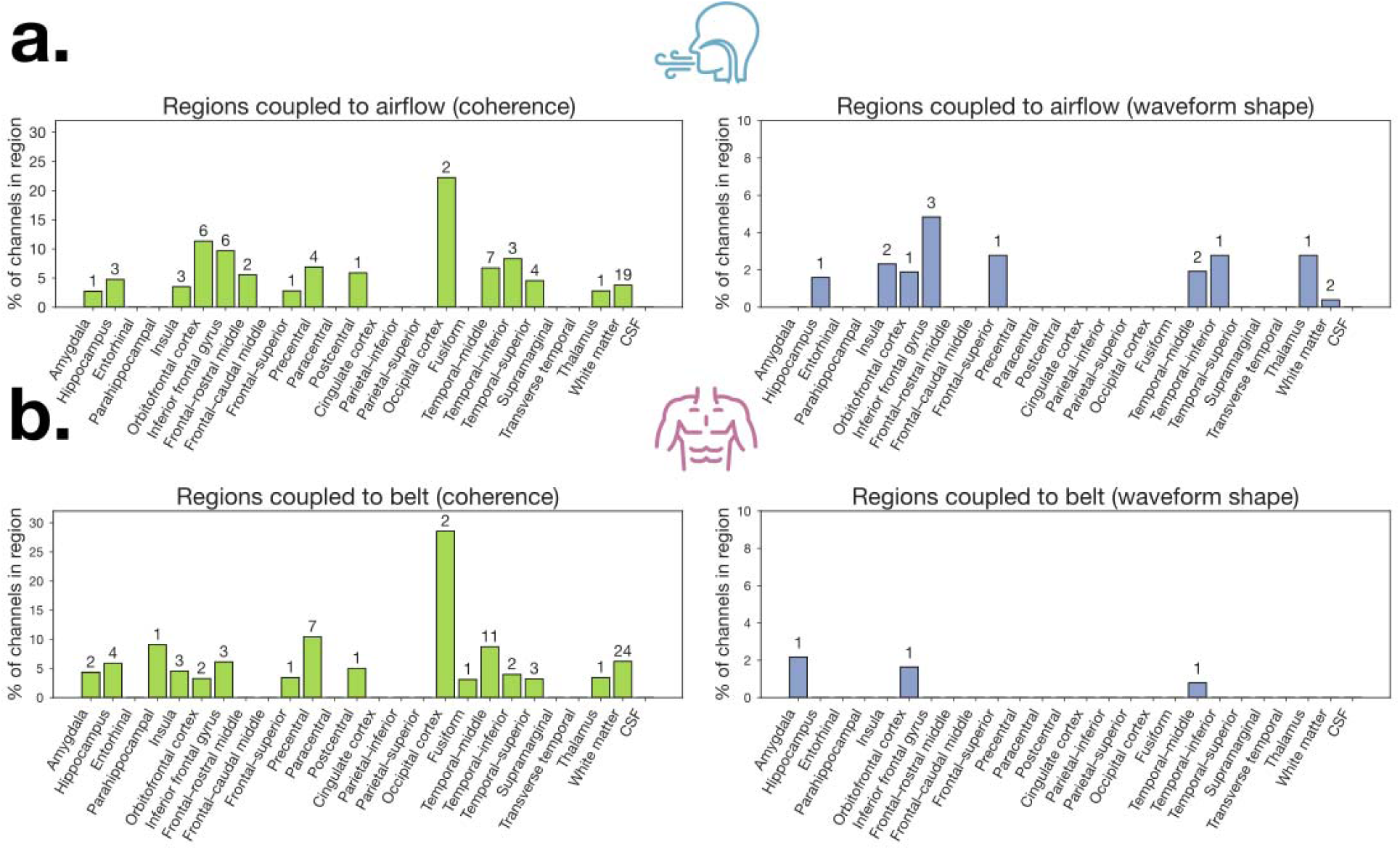
Distribution of electrodes across brain regions showing respiration-brain coupling for airflow and belt signals. **a.** Airflow-based coupling. Left: Percentage of total channels per region exhibiting significant coherence with nasal airflow (green). Right: Percentage of total channels per region showing significant respiration-locked waveform shape (blue). **b.** Belt-based coupling. Left: Percentage of total channels per region exhibiting significant coherence with the respiratory belt signal (green). Right: Percentage of total channels per region showing significant respiration-locked waveform shape (blue). Numbers above bars represent raw counts of channels.

Significant coherence with airflow was observed across distributed limbic and cortical regions (**Fig. 3a**, left). Although occipital cortex showed the highest proportion of coherent channels, this reflects only 2 significant channels out of 9 total and should be interpreted cautiously. Among regions with broader coverage, the most prominent coherence was observed in orbitofrontal cortex, inferior frontal gyrus, temporal-inferior, precentral, temporal-middle, and postcentral cortex, with additional coupling observed in hippocampus, amygdala, and insula. Belt coherence showed a similarly distributed pattern (**Fig. 3b**, left), again led nominally by occipital cortex (2/7 channels), with more robustly represented coupling in precentral gyrus, parahippocampal cortex, temporal-middle, and inferior frontal gyrus, with additional contributions from hippocampus, amygdala, and insula. However, visual inspection revealed that coherence alone did not necessarily imply clear cycle-by-cycle alignment in the time domain (**Fig. 2c**, middle row). Thus, additional screening was performed to ensure waveform-level correspondence.

Following coherence selection, we applied two additional criteria: cross-correlation within respiration-centered windows and a respiration PMI. Cross-correlation analyses demonstrated that a subset of coherent channels exhibited coupling with consistent temporal lag structure (**Fig. 2c**). These sequential filters reduced the dataset to: 18 airflow channels (across 7 participants) exhibiting significant coherence, cross-correlation, and PMI, and 4 belt channels (across 3 participants) passing all screening criteria.

Among airflow-coherent channels, waveform shape modulation was concentrated in a smaller subset of regions (**Fig. 3a**, right), with highest proportions in inferior frontal gyrus, thalamus, frontal-superior cortex, and temporal-inferior cortex, followed by insula, lateral orbitofrontal cortex, temporal-middle cortex, and hippocampus. In contrast, waveform shape modulation for belt signals was substantially sparser (**Fig. 3b**, right), with only isolated contributions from the amygdala, lateral orbitofrontal cortex, and temporal-middle cortex. Among airflow-coupled channels passing all waveform shape criteria, two channels localized to white matter. While these channels passed all screening criteria, white matter channels are not expected to reflect local neural generators and likely reflect volume conduction from adjacent gray matter sources.

### Respiratory waveform shape is coupled to neural waveform shape

We next asked whether cycle-by-cycle variation in respiration morphology relates to neural oscillation morphology. Using Bayesian linear mixed-effects models, neural waveform features were modeled as a function of matched respiratory features using a within-between decomposition. All features were scaled by their pooled (grand) standard deviation prior to modeling, such that standardized slopes (β) reflect within-subject coupling strength and are directly comparable across features.

Airflow waveform features showed consistent positive relationships with corresponding neural waveform features (**Fig. 4a–b**): Rise time: β = 0.17 [95% CrI: 0, 0.39], P(β > 0) = 0.97; Decay time: β = 0.13 [0.02, 0.23], P(β > 0) = 0.99; Cycle duration: β = 0.19 [0.07, 0.33], P(β > 0) = 1; Inhale peak AUC: β = 0.16 [0.07, 0.26], P(β > 0) = 1.00; Exhale trough AUC: β = 0.12 [0.03, 0.23], P(β > 0) = 0.99. Sharpness and symmetry features were inconclusive: Inhale peak sharpness: β = −0.01 [−0.06, 0.03], P(β > 0) = 0.32; Exhale trough sharpness: β = 0.02 [−0.02, 0.07], P(β > 0) = 0.87; Rise-decay symmetry: β = 0.07 [−0.09, 0.23], P(β > 0) = 0.80. Additionally, for features showing strong posterior probability for coupling, the real model posteriors were visibly displaced from the null model posteriors fit on cycle-shuffled data, while sharpness and symmetry posterior distributions overlapped substantially with the null (**Fig. 4a**). These results indicate strong support for positive relationships between airflow morphology and neural morphology for temporal and amplitude-related features, but not for sharpness or symmetry.

**Fig. 4.**
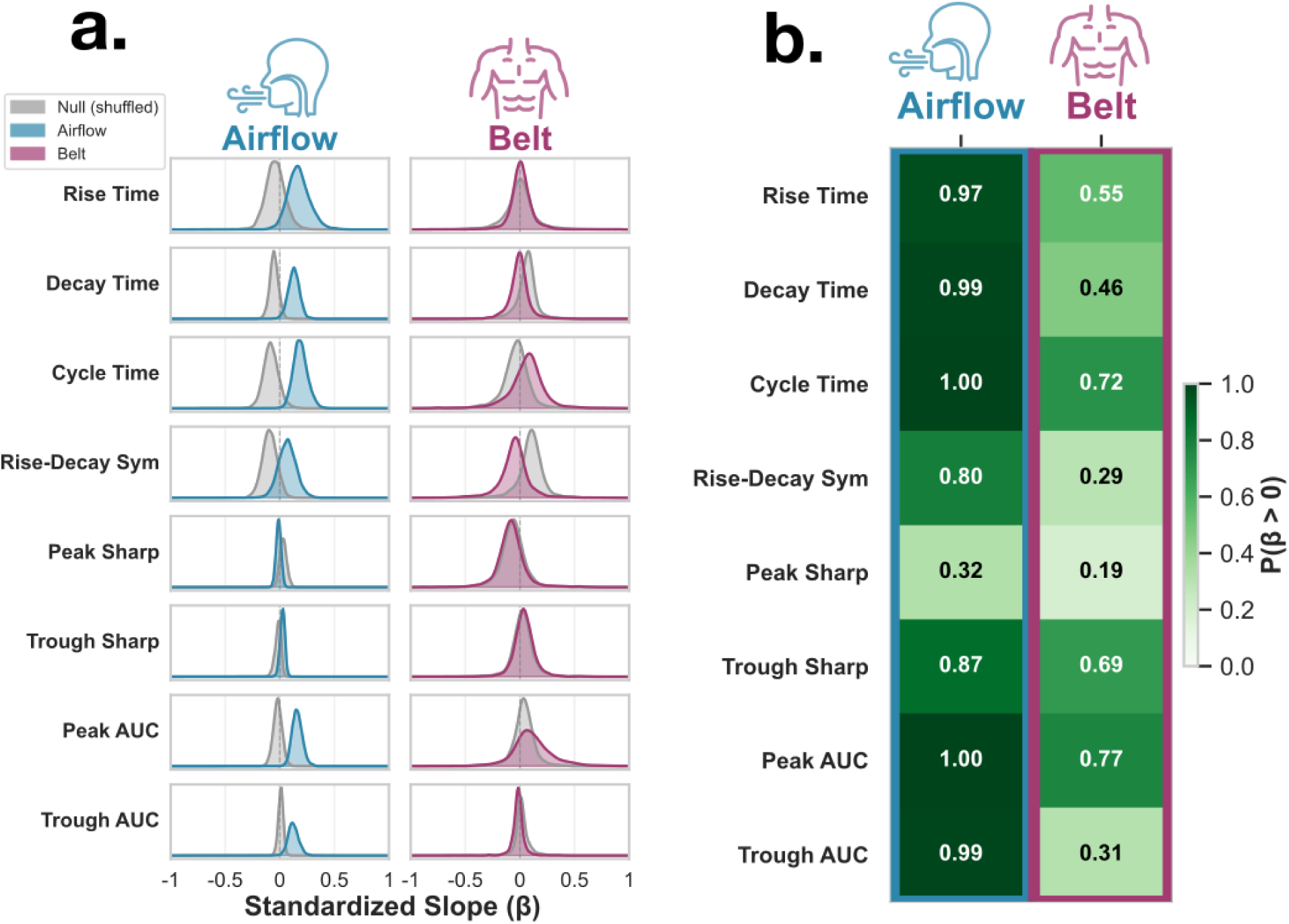
Respiratory waveform shape features are related to neural waveform shape features. **a.** Posterior distributions of standardized slopes (β) from Bayesian linear mixed effects models relating respiratory features (rise time, decay time, cycle duration, rise-decay symmetry, inhale peak sharpness, exhale trough sharpness, inhale peak AUC, exhale trough AUC) to corresponding neural features. Slopes are standardized (z-scored predictors and outcomes), such that coefficients reflect effect sizes in SD units. Blue and purple distributions represent the posterior densities for airflow and belt, respectively. Gray distributions represent null model posteriors fit on data in which the within-subject respiratory predictor was shuffled across cycles to break cycle-by-cycle coupling. Positive values indicate that increases in respiratory feature magnitude are associated with increases in the corresponding neural feature. **b.** Posterior probability that each standardized slope is positive, P(β > 0), for airflow and belt models. Darker shading indicates stronger evidence for a positive relationship. Airflow demonstrates high posterior support for positive associations for rise time, decay time, cycle duration, inhale peak AUC, and exhale trough AUC (P ≥ 0.97), whereas belt shows more moderate or uncertain evidence across features.

Belt-derived results were uniformly inconclusive across all waveform features: Rise time: β = 0.01 [−0.18, 0.23], P(β > 0) = 0.55; Decay time: β = −0.01 [−0.21, 0.19], P(β > 0) = 0.46; Cycle duration: β = 0.06 [−0.34, 0.38], P(β > 0) = 0.72; Rise-decay symmetry: β = −0.05 [−0.38, 0.22], P(β > 0) = 0.29; Inhale peak AUC: β = 0.11 [−0.26, 0.55], P(β > 0) = 0.77; Exhale trough AUC: β = −0.02 [−0.14, 0.10], P(β > 0) = 0.31; Inhale peak sharpness: β = −0.08 [−0.35, 0.23], P(β > 0) = 0.19; Exhale trough sharpness: β = 0.04 [−0.15, 0.25], P(β > 0) = 0.69. Correspondingly, the belt model posterior distributions were largely overlapping with the null model posteriors across all features (**Fig. 4a**). Thus, while airflow features consistently showed evidence for coupling with neural waveform morphology, belt signals did not.

The cycle-by-cycle scatter plots illustrate this pattern visually: airflow regression lines show mostly consistent positive slopes across channels for temporal features (rise time, decay time, cycle duration) and amplitude features (inhale peak and exhale trough AUC), whereas belt regression lines are variable in direction, and sharpness slopes are flat in both modalities (**Fig. 5**).

**Fig. 5.**
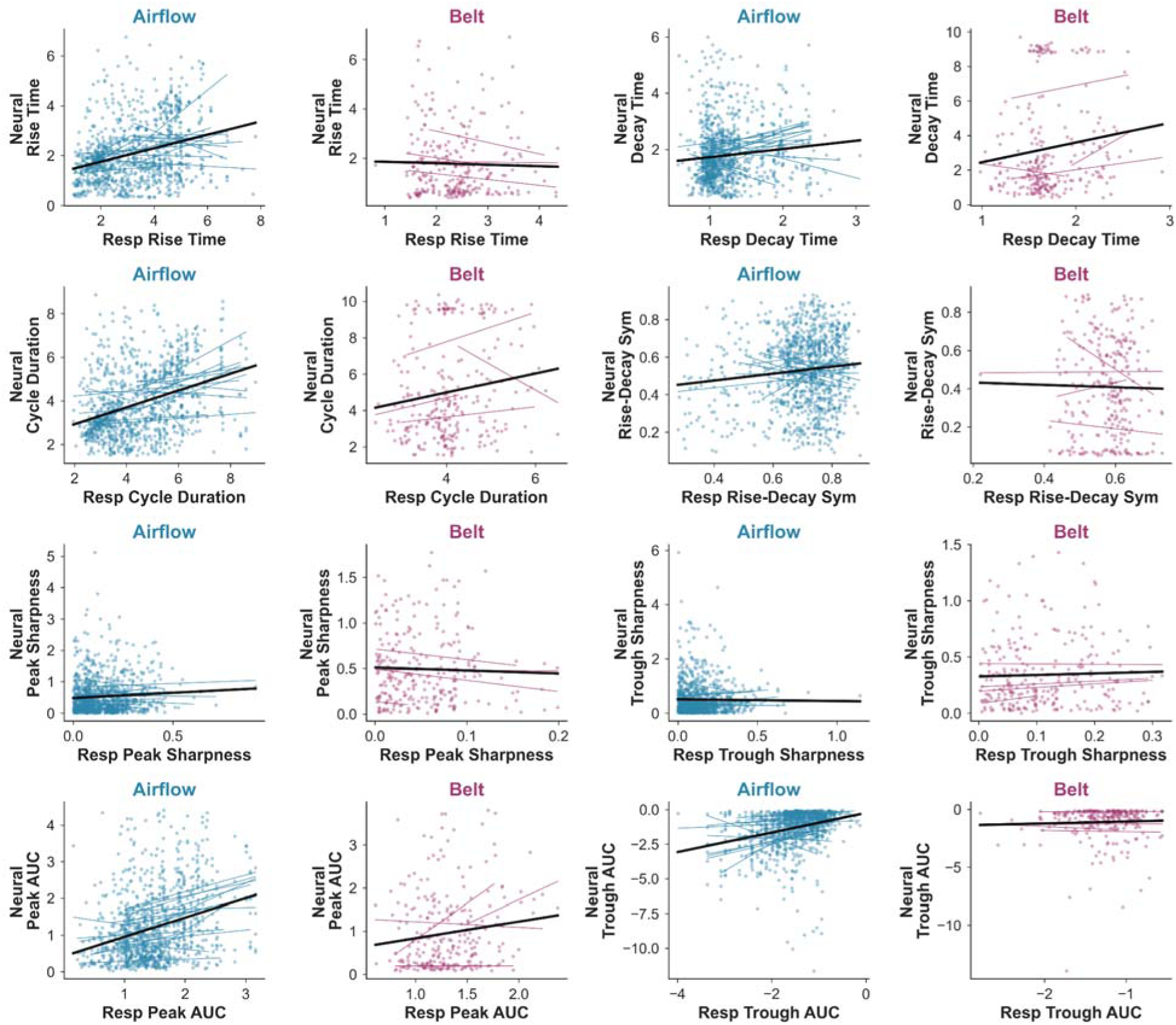
Cycle-by-cycle scatter plots of respiratory and neural waveform shape features. Each point represents a single matched respiratory-neural cycle. Lines indicate within-channel regression fits; black lines indicate the overall regression fit across all channels and subjects. Airflow (blue) and belt (purple) modalities are shown separately for each feature pair: rise time, decay time, cycle duration, rise-decay symmetry, inhale peak sharpness, exhale trough sharpness, inhale peak AUC, and exhale trough AUC. Positive slopes across participants are most consistent for airflow-derived temporal features (rise time, decay time, cycle duration) and amplitude-related features (inhale peak and exhale trough AUC), reflecting the within-subject coupling captured by the Bayesian models. Sharpness features show flatter and more variable slopes in both modalities. Belt regression lines are more inconsistent in direction across channels.

## Discussion

Across three independent human sEEG datasets, we confirm widespread respiratory-neural coherence across limbic and cortical regions. Among coherent channels, a subset also exhibit morphological coupling such that respiratory waveform shape is related to neural oscillation waveform shape on a cycle-by-cycle basis, an effect substantially stronger for nasal airflow than belt recordings.

Humans breathe in asymmetric, nonsinusoidal shapes, with varying inhalation and exhalation durations, pauses between breaths, and differences in the volume/intensity of air that gets inhaled and exhaled. Prior work has demonstrated that there are rhythms in the brain that are statistically coherent with breathing (Tort et al., 2025); however, this metric averages over cycle-level variability and doesn’t imply that the neural oscillations are systematically synchronized with the precise waveform shapes of the respiration. We therefore isolated neural channels with waveform coupling to both airflow and belt waveforms, matched these signals on a cycle-by-cycle basis, parameterized their waveform shapes, and found that respiration shape is reliably related to neural shape in expected regions.

This relationship suggests that respiration may modulate the shapes of neural waveforms in a temporally-precise manner. While not causal proof, this cycle-by-cycle relationship implies a tight coupling between the peripheral and central nervous system in a manner that, until now, has never been demonstrated. Neural oscillation waveform shape is thought to represent properties of the underlying physiological generators of the local field potential, including the synchrony of various postsynaptic potentials and the timing of population-level firing (S. R. Cole & Voytek, 2017; Trimper et al., 2014). If respiration shapes the waveform morphology of neural oscillations on a cycle-by-cycle basis, as we demonstrate here, this implies that breath-to-breath variation may directly modulate the temporal structure of neural activity in limbic and cortical regions. In other words, the asymmetry of the breath cycle – how quickly or slowly air flows in and out – may impact the duration of neural up- and down-states with downstream consequences for the time windows available for spike probability and inter-regional communication (Karalis & Sirota, 2022).

The substantially stronger coupling observed for nasal airflow compared to the respiratory belt is consistent with the interpretation that the underlying driver of this effect arises from the nasal airflow pathway and its early projections into limbic circuitry rather than from the mechanical act of breathing itself (Fontanini et al., 2003; Juventin et al., 2022; Lockmann et al., 2016; Zelano et al., 2016). However, recent work has demonstrated that forebrain synchronization with breathing in humans persists even when airflow bypasses the oronasal passages entirely (Dlouhy et al., 2025). The weaker coupling from the belt could therefore reflect not the absence of a mechanistic link, but rather the limited fidelity of the belt as a recording instrument, which coarsely measures chest wall displacement rather than nasal airflow dynamics directly (Johnson et al., 2006). Nasal airflow recordings preserve certain temporal features of the breath cycle that the belt signal misses, such as the post-expiratory pause (PEP). This waveform feature carries functional significance for limbic circuits and predicts depression and life satisfaction (Diez et al., 2025).

The regional distribution of waveform-coupled channels is broadly consistent with known anatomy of respiration-brain pathways. Among airflow-coupled channels passing all screening criteria, the largest concentrations were observed in inferior frontal gyrus (3 channels), insula (2 channels), and thalamus (1 channel), with isolated contributions from frontal-superior and temporal-inferior cortex. Belt-coupled channels were even sparser, with single channels in amygdala, insula, and fusiform cortex. A direct anatomical pathway links nasal airflow to limbic and cortical regions: mechanoreceptors in the nasal epithelium project via the olfactory bulb to piriform cortex and from there to the amygdala, hippocampus, and insula, providing a substrate for airflow-specific entrainment of forebrain activity (Fontanini et al., 2003; Juventin et al., 2022; Zelano et al., 2016). The insula is known to be strongly synchronized to breathing, particularly anterior insula (Assi et al., 2025; Dlouhy et al., 2025) due to its role as an interoceptive hub integrating ascending visceral signals to predict bodily states ((Bud) Craig, 2009; Chen et al., 2021; Khalsa et al., 2009). Temporal and frontal regions are likewise consistent with broad forebrain distribution of respiration-coherent channels during wakefulness (Dlouhy et al., 2025), and resting-state fMRI work has specifically shown that nasal – but not oral – breathing promotes functional connectivity between olfactory regions and lateral frontal cortex including IFG, consistent with our finding that IFG waveform coupling was observed for airflow but not belt signals (Mohammadi et al., 2025). Furthermore, the concentration of waveform-coupled channels in insula-adjacent limbic and cortical regions is consistent with their role as core nodes of the salience network, which integrates interoceptive and affective signals, and is linked to precise temporal features of airflow (Diez et al., 2025). Additionally, recent work has demonstrated that thalamus is an important region for respiratory information; single-unit recordings in human thalamus were shown to actively encode the temporal structure and phases of the breath, preferentially spiking during inhalation versus exhalation (De Falco et al., 2024). Finally, two airflow-coupled channels were localized to white matter, which is unlikely to reflect local synaptic activity and most plausibly arises via volume conduction from nearby gray matter sources (Herrero et al., 2018). Notably, the overwhelming concentration of waveform-coupled channels in gray matter and the relative rarity of white matter channels is itself consistent with the interpretation that the observed coupling reflects genuine neural activity rather than a respiratory mechanical artifact.

There are a number of limitations to the present study. Our method isolated only a small number of channels exhibiting significant waveform shape coupling, particularly for belt recordings, reflecting the difficulty of detecting this signal even when coherence is present. Notably, only four channels across three participants passed all screening criteria for belt-based waveform shape analysis, making the belt models severely underpowered and their null results difficult to interpret in isolation. That said, the fact that belt channels were disproportionately filtered out by the time-domain screening steps, despite comparable numbers of coherent channels, is itself informative, even if the statistical models were underpowered. Additionally, all participants were epilepsy patients undergoing invasive monitoring, with electrode placement determined by clinical need rather than experimental design. The underlying pathology, antiseizure drug regimens, and uneven cortical and subcortical coverage across participants may limit generalizability to healthy populations (see **Table 1** for information on participants).

Future research should leverage volitional breath control to determine whether controlling respiratory waveform shape influences the shape of neural waveforms in a causal manner. Unlike passive breathing at rest, volitional breathing recruits frontal, motor, and premotor circuits (Herrero et al., 2018), introducing a top-down component to the respiration-brain pathway. If deliberate manipulation of breath shape drives corresponding changes in the neural waveform, this would suggest a bidirectional pathway in which frontal control of breathing feeds back onto the limbic and cortical dynamics that breathing entrains. Beyond controlled manipulations of breathing, future work could determine whether respiratory waveform shape features – not just respiratory phase – predict trial-by-trial variability during cognitive, affective, or perceptual tasks. This would help establish whether the cycle-by-cycle morphological coupling we demonstrate here has direct behavioral implications.

## Data availability

All code used for all analyses and plots are publicly available on GitHub at https://github.com/voytekresearch.

## Acknowledgements

This work was supported by the NIH National Institute of Mental Health grant R61MH135109. We are grateful to members of the Voytek Lab for their advice and feedback on the manuscript.

## Author contributions

E.K.R. and B.V. conceived of the analysis framework. E.K.R. wrote the analysis code and analyzed the data. G.Z., A.S., J.M.R., S.U.S., C.O.O., S.A.T., M.K., M.R.M., A.E.R., S.K., C.Z., and B.D. provided data and advised analyses. E.K.R. and B.V. wrote the manuscript, and all authors edited the manuscript.

Competing interests the authors declare no competing interests.

---

**Fig. S1.**
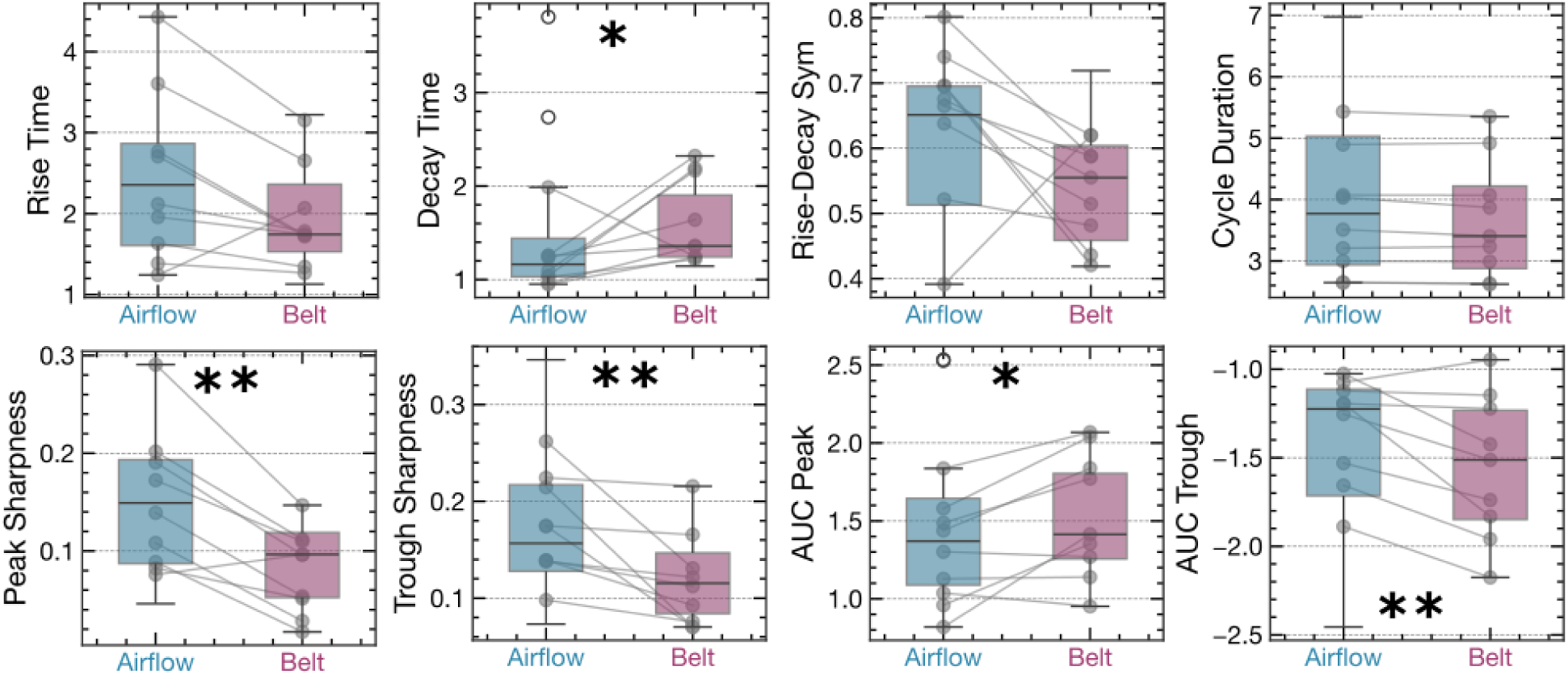
Respiration waveform shape quantified across modalities. Boxplots show the distribution of median per-participant waveform shape features for airflow (blue) and belt (purple) recordings for participants with both modalities (N = 10). Each point represents one participant; gray lines connect paired observations from the same participant (n = 10 participants with both modalities). Asterisks indicate significant paired differences by Wilcoxon signed-rank test (* p < 0.05, ** p < 0.01).

